# MspA Porin as a Local Nanopore Probe for Membrane-bound Proteins

**DOI:** 10.1101/2021.10.15.464579

**Authors:** David P. Hoogerheide, Philip A. Gurnev, Jens Gundlach, Andrew Laszlo, Tatiana K. Rostovtseva, Sergey M. Bezrukov

**Affiliations:** Center for Neutron Research, National Institute of Standards and Technology, Gaithersburg, MD 20899; Section on Molecular Transport, Eunice Kennedy Shriver National Institute of Child Health and Human Development, National Institutes of Health, Bethesda, MD 20892; Department of Physics, University of Washington, Seattle, WA 98105

## Abstract

Nanopore sensing is based on detection and analysis of nanopore transient conductance changes induced by analyte capture. We have recently shown that α-Synuclein (αSyn), an intrinsically disordered, membrane-active, neuronal protein implicated in Parkinson disease, can be reversibly captured by the VDAC nanopore. The capture process is a highly voltage dependent complexation of the two proteins where transmembrane potential drives the polyanionic C-terminal domain of αSyn into VDAC—exactly the mechanism by which generic nanopore-based interrogation of proteins and polynucleotides proceeds. The complex formation, and the motion of αSyn in the nanopore, thus may be expected to be only indirectly dependent on the pore identity. Here, we confirm this prediction by demonstrating that when VDAC is replaced with a different transmembrane pore, the engineered mycobacterial porin M2MspA, all the qualitative features of the αSyn/nanopore interaction are preserved. The rate of αSyn capture by M2MspA rises exponentially with the applied field, while the residence time displays a crossover behavior, indicating that at voltages >50 mV M2MspA-bound αSyn largely undergoes translocation to the other side of the membrane. The translocation is directly confirmed using the selectivity tag method, in which the polyanionic C-terminal and neutral N-terminal regions of αSyn alter the selectivity of the M2MspA channel differently, allowing direct discrimination of translocation vs retraction for single αSyn molecules. We thus prove that the physical model of the motion of disordered protein chains in the nanopore confinement and the selectivity tag technique are not limited to VDAC but are broadly applicable to nanopore-based protein detection, analysis, and separation technologies.

## Introduction

Among different single-molecule sensing techniques, sensing with nanopores stands out as one of the most versatile and rapidly evolving. The technique is based on the capture of charged and neutral biomolecules into the nanopore confinement, followed by analysis of the resulting changes in the current of small ions through the nanopore driven by an applied transmembrane potential. Due to its nanoscale dimensions, a hole in a thin membrane separating two reservoirs filled with electrolyte solution allows, in direct analogy to the Coulter-counting principle, detection and examination of “particles” of molecular size (1). In the case of charged nanopores, a finite selectivity between cations and anions arises, so that an extra driving force can result from the difference in electrolyte concentration at the opposite sides of the membrane. The function of these driving forces is not limited to generation of small-ion current; in many cases they are also responsible for the capture, retention, and translocation of the captured molecules. Time-resolved changes in the ionic current report on the charge, structure, and dynamics of the molecules within the nanopore (2, 3), post-translational and other single-residue modifications (4-6), and protein-ligand interactions (7). There is a significant current effort to develop nanopore-based protein sequencing and proteomic analysis (8-10).

While a nanopore sensing system can be constructed entirely from biological molecules—lipids comprise the membrane, whereas biological ion channels make excellent nanopores—it has only recently become clear that voltage-induced capture of polymeric, polyvalent analytes may itself have physiological relevance. The mitochondrial outer membrane voltage-dependent anion channel (VDAC) forms a transmembrane voltage-activated complex with two known peripheral membrane proteins, dimeric tubulin and αSyn (11). Assembly of the voltage-activated complex proceeds completely analogously to capture of a polynucleotide in a nanopore: under the influence of the transmembrane potential, the disordered, polyanionic C-terminal domains of membrane-bound tubulin or αSyn are threaded through VDAC until arrested by a membrane anchor. This metastable complex eventually disassembles either by retraction of the C-terminal polypeptide from VDAC, or, in the case of the intrinsically disordered αSyn, membrane anchor detachment and translocation of the entire protein (12-14).

The evidence for voltage-activated complexation of VDAC with tubulin and αSyn is largely derived from *in vitro* experiments in which individual VDAC molecules are reconstituted into a biomimetic planar lipid bilayer. These experiments proceed exactly as the nanopore sensing ones: the ionic current through VDAC is monitored in the presence of tubulin or αSyn; the voltage-dependent rates of onset and recovery (“on-rate” and “off-rate”) of transient current reductions induced by these molecules are then determined. These voltage-dependent rates can be quantitatively described by a free energy profile incorporating the physical properties of the analyte polypeptides, rather than those of VDAC (12, 15, 16). In the case of αSyn, which can translocate the membrane through VDAC, the free energy profile accurately predicts the translocation probability (13, 14). Together, this constitutes strong evidence that the mechanism of the observed interaction is voltage-induced insertion of the C-terminal domains into the VDAC nanopore.

An alternate interpretation is gating of the VDAC channel induced by tubulin or αSyn. VDAC is best known for its eponymous property of voltage-induced gating (17, 18): under sufficiently high transmembrane potential (typically 30 mV or larger, depending on lipid composition (19- 21), ionic strength (19, 20), or pH (22, 23)), VDAC spontaneously transitions into relatively low-conductance states. The conductance of the gated state varies but is similar to that observed from interaction with tubulin or αSyn, while the duration of the gated state is typically orders of magnitude longer than the reductions in conductance induced by tubulin or αSyn. Thus, the question remains whether the interaction with tubulin or αSyn might involve an alternate mechanism by which tubulin or αSyn induce short-lived gated states without passage of the C-terminal domain through the VDAC pore. In this case the close correspondence of the measured voltage-dependent rates and translocation probability to the free energy landscape modeling predictions would be an uncanny coincidence. In addition, the anion selectivity of the VDAC channel would be expected to be of critical importance in binding the polyanionic domains of tubulin or αSyn, as these domains have been shown to be essential for observing conductance modulations (24).

To distinguish between these two interpretations, the conclusive experiment is to replace the VDAC channel with a channel that is neither anion selective nor gates. Here, we report the results of just such an experiment, in which VDAC is replaced with a variant of a widely used proteinaceous mycobacterial nanopore, M2MspA (hereafter “MspA”). The nanopore is modified from wild type MspA to eliminate negatively charged residues from the constriction (D90N/D91N/D93N) and add positively charged residues to the channel walls (D118R/E139K/D134R) (25, 26). Removal of the charged residues from the pore constriction eliminates its ion selectivity. We show that all the characteristic features of the VDAC-αSyn complexation are preserved with the M2MspA pore, thus conclusively establishing insertion of αSyn’s C-terminal domain into the pore interior as the complexation mechanism.

## Materials and methods

### Experiment

Mycobacterium smegmatis porin A is modified from the wild type via D90N/D91N/D93N/D118R/E139K/D134R mutations to enable nanopore DNA sequencing (25). A diphytanoylphosphatidylcholine (DPhPC) bilayer was formed from lipid monolayers according to the method of Montal and Mueller (27). MspA was added to the *cis* side of the membrane in the amount of 10 pM to 50 pM (M = mol/L) in 0.1 % octyl-polyoxyethylene; the opposite side is referred to as the *trans* side (Figure 1). Individual channels had a conductance of about 2 nS in 1 M KCl at 100 mV. In this manuscript, voltage polarities are defined as positive when the *cis* side has the higher potential. To create a salt concentration gradient, the *cis* side was perfused to a final concentration of 0.2 M KCl, while the *trans* side remained at 1.0 M KCl, both sides buffered with 5 mM HEPES at pH 7.4. Junction potentials with the Ag/AgCl electrodes were minimized by connecting the electrodes with 2.0 M KCl / 2% agarose bridges. Recombinant αSyn, a kind gift from Dr. Jennifer Lee (NHLBI, NIH), was added to the *trans* side at 50 nM concentrations. Current recordings at each voltage were collected by an Axopatch 200B amplifier (Molecular Devices, Sunnydale, CA) with a 4 μs sampling interval, hardware filtered by a 10 kHz inline 8-pole Bessel filter (9002, Frequency Devices, Ottawa, IL), and directly saved into computer memory with Clampex 10 software (Molecular Devices). All experiments were performed at room temperature (22 °C ± 2 °C).

**Figure 1.**
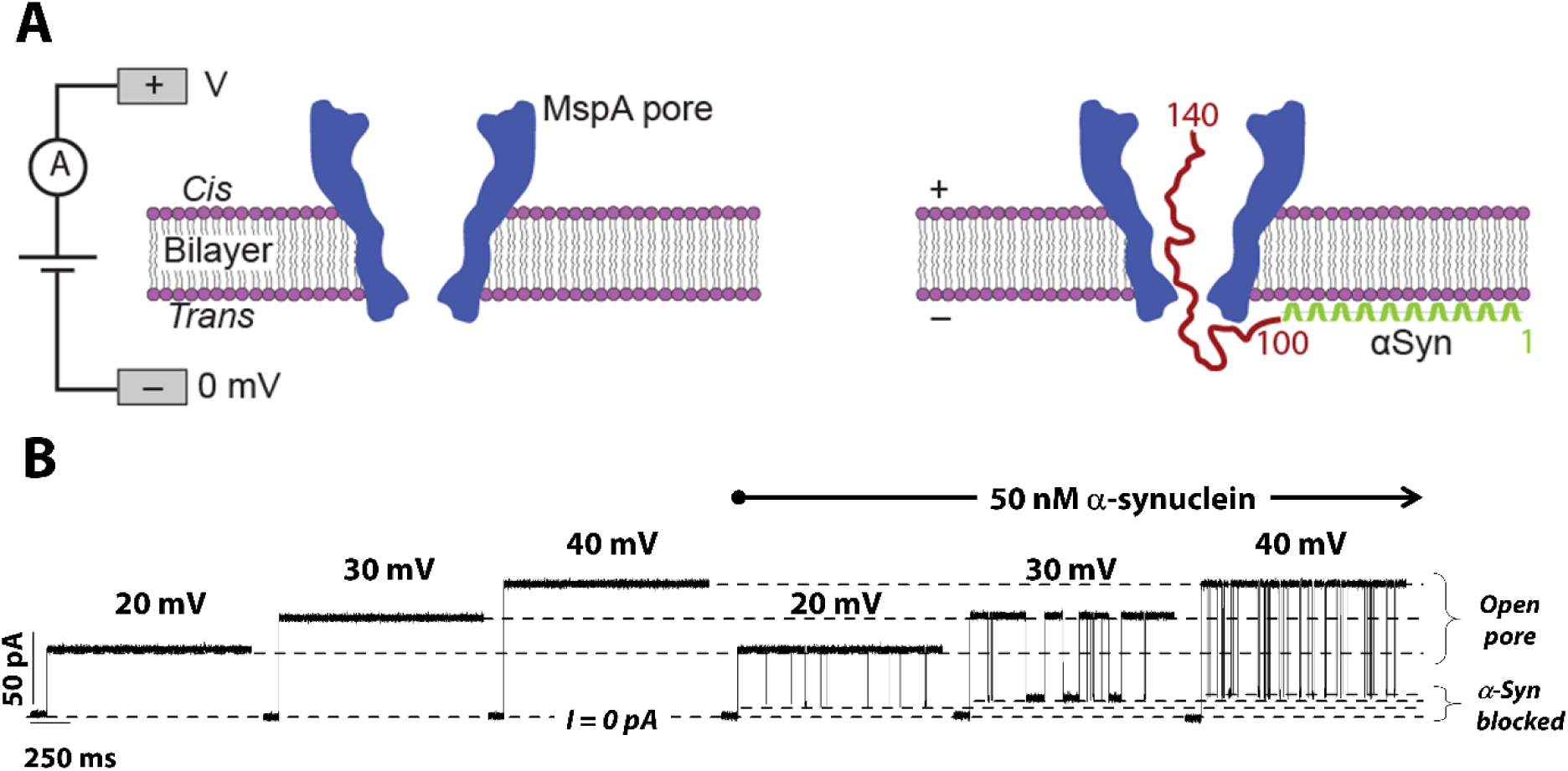
(A) Schematic of the experimental setup with reconstituted MspA in a DPhPC bilayer before (left) and after (right) addition of 50 nM αSyn to the *trans*-side of the planar lipid bilayer. (B) Representative traces of the current through a single MspA nanopore before (left) and after (right) αSyn addition, at different applied voltages (positive relative to the side of αSyn addition). The bilayer was bathed with 1.0 M KCl, 5 mM HEPES, pH 7.4 solutions on both sides of the membrane. αSyn induces time-resolved rapid blockages of the small-ion current passing through the MspA nanopore.

### Data analysis

For single-channel data analysis by Clampfit 10.3, a digital 8-pole Bessel low-pass filter set at 5 kHz or 2 kHz was applied to current recordings, and then, individual events of current blockages were discriminated and kinetic parameters were acquired by fitting single exponentials to logarithmically binned histograms (28) as described previously (24, 29). Each experiment was replicated 3 times; the results of one experiment are shown for clarity.

## Results

### αSyn/MspA interaction in symmetrical salt conditions

MspA was reconstituted into planar bilayers exposed to 1 M KCl buffer on each side. The octameric MspA channels are inserted in a uniform direction, judging by the reproducible asymmetry of the voltage dependence of the observed pore’s conductance: at positive voltages in the 30 mV to 90 mV range, pores pass ∼ 20 % less current than at negative voltages of the same magnitudes. It was shown earlier (30) that this conductance asymmetry reflects MspA’s tendency to reconstitute in bilayers with its large extracellular opening facing the side of the protein’s addition to the bilayers (*cis*-side in our convention).

In the absence of αSyn, MspA produces steadily conductive pores with few spontaneous stepwise current fluctuations up to an absolute applied voltage of 150 mV. When αSyn (50 nM) was added to the *trans*-compartment of the chamber, i.e. to the side of the membrane where the shorter and narrower MspA pore opening was located, it produced characteristic, transient, stepwise reductions in the current to about 20 % of the original pore conductance (Fig. 1). These current blockages, or “events”, were only observed at the voltage polarities positive from the side opposite to that of αSyn addition; no events were observed with voltages of inverted polarity or at either polarity with *cis*-side-only αSyn additions.

These observations suggest that the mechanism of αSyn/MspA interaction is essentially the same as that observed earlier with the α-hemolysin and VDAC pores (Gurnev, Yap et al. 2014, Rostovtseva, Gurnev et al. 2015, Hoogerheide, Rostovtseva et al. 2021): membrane-bound αSyn is drawn into the pore by the polyanionic C-terminal domain. Similarly, blockages of MspA by αSyn are highly voltage-dependent; Fig. 2 summarizes the kinetics of this interaction.

**Figure 2.**
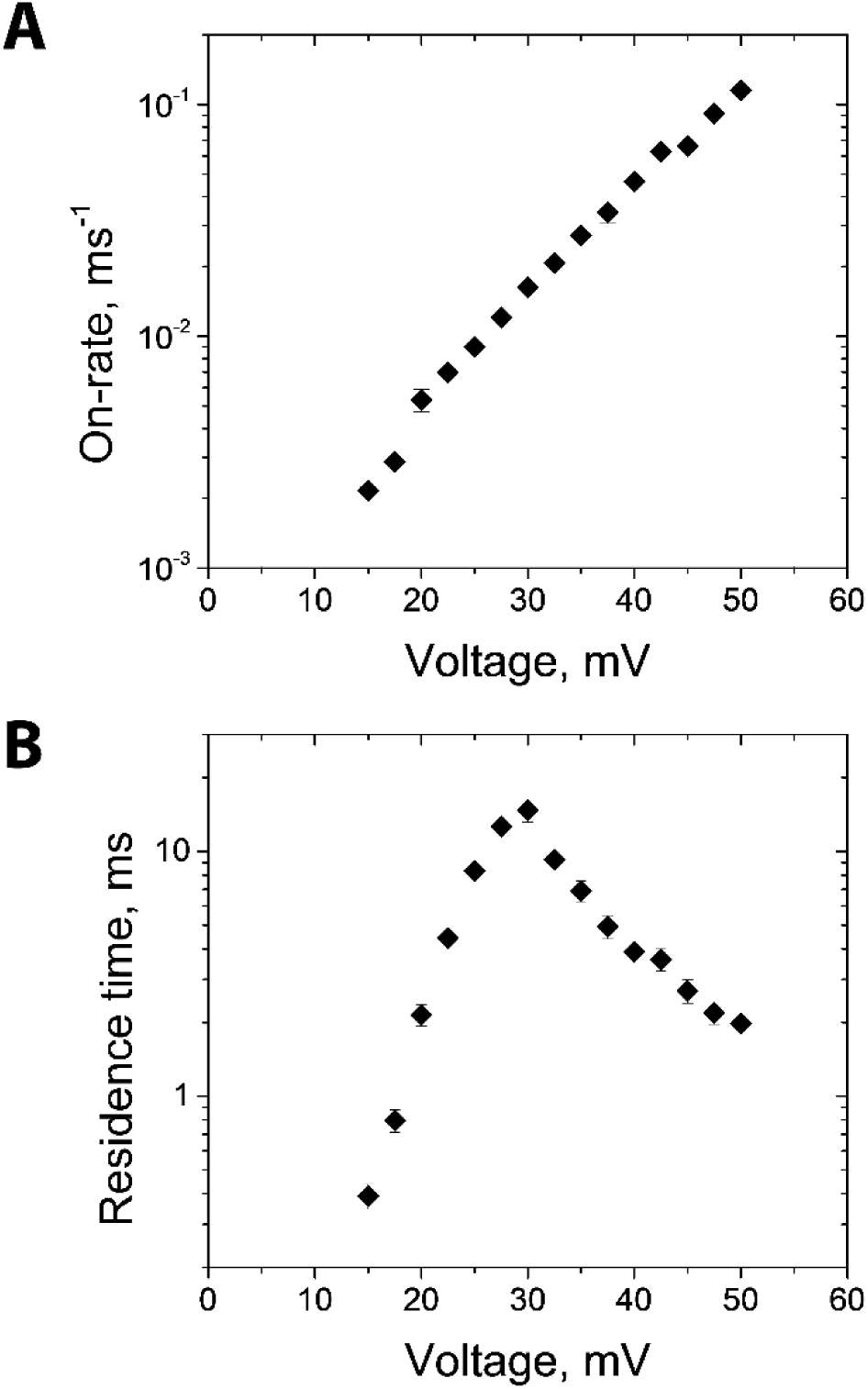
αSynuclein blocks the MspA pore in a highly voltage-dependent manner. (A) Capture rate of αSyn into MspA increases exponentially with absolute applied voltage. (B) The blockage duration has a biphasic dependence on the applied voltage (exponential growth in the regimes of 15 mV to 30 mV and a decrease above 30 mV). Error bars represent 68% confidence intervals.

The distribution of times between blockage events when the MspA pore is unblocked, τ_on_, is well described by a single exponential function at all the studied voltages, and the effective on-rate of this interaction (<τ_on_>^-1^) grows exponentially with the applied voltage (Fig. 2A).

The dependence of residence time, τ_off_, is biphasic in the applied voltage (Fig. 2B). The exponential increase of *τ_off_* at absolute voltages <30 mV has been shown in the VDAC channel to arise from a deepening potential well as the C-terminal domain is subjected to stronger electrical forces that prevent escape. At higher applied voltages, the *τ_off_* dependence changes, and *τ_off_* values begin to decrease with voltage. With VDAC, this has been unequivocally shown to be due to translocation of αSyn through the channel. In the following section, it will be demonstrated for MspA.

### Voltage-dependent translocation probability, asymmetrical salt conditions

We have shown earlier with membrane-reconstituted mitochondrial VDAC that the translocation success of αSyn through this nanopore can be monitored with precise resolution by imposing an electrolyte (salt) gradient across the membrane (12, 14, 16). The technique takes advantage of the fact that the presence of a charged polypeptide chain alters a nanopore’s ion selectivity, and under salt gradient conditions this translates to a shift in the reversal potential and hence in the ionic current. Serendipitously, αSyn is well suited for exploitation of this effect. The primary structure of the 140-residue αSyn comprises two regions of very different average charge density: the first 97 residues’ total charge is nearly neutral (+3*e*), while the remaining C-terminal 43 amino acids carry a net charge of –15*e*. The boundary between these two regions thus acts as a fiduciary point, and its passage through the pore is marked by transition of the ionic current between two sublevels corresponding to the presence of the two regions of αSyn in the nanopore constriction.

Here, we demonstrate this effect using MspA. Upon perfusion of the *cis*-compartment of the chamber to obtain a final KCl concentration of 0.2 M (compared to the original 1.0 M), the current blockages produced by αSyn in MspA are getting split into two readily distinguishable current states within a single blockage event (Fig. 3A). To establish the characteristic reversal potentials, and hence the ion selectivity associated with them, these two substates of the blocked conductance are observed over a range of applied voltages (Fig. 3B). From the current-voltage relationships, shown on this graph, we indeed confirm that one of the αSyn-blocked states of the MspA pore is cation-selective (reversal potential is 23.2 ± 0.7 mV, which corresponds to a 0.23 Cl^−^ / 1 K^+^ ratio; error bars represent 68% confidence intervals estimated from linear regression) and the other one is essentially non-selective (reversal potential is equal to −3.7 ± 0.6 mV, a 1.2 Cl^−^/1 K^+^ ratio). By contrast, the open state of the pore is non-selective, characterized by a reversal potential close to zero. These values allow straightforward assignment of the two blocked states to the two regions of αSyn: the negatively charged C-terminal produces the deeper blocking cation-selective state and the N-terminal domain (or, more likely, the cluster of positive charge from residues 96 to 102) produces the shallower, non-selective one. The observed fluctuations in ion selectivity of the blocked states of MspA report on the motion of the αSyn fiduciary boundary through the MspA constriction.

**Figure 3.**
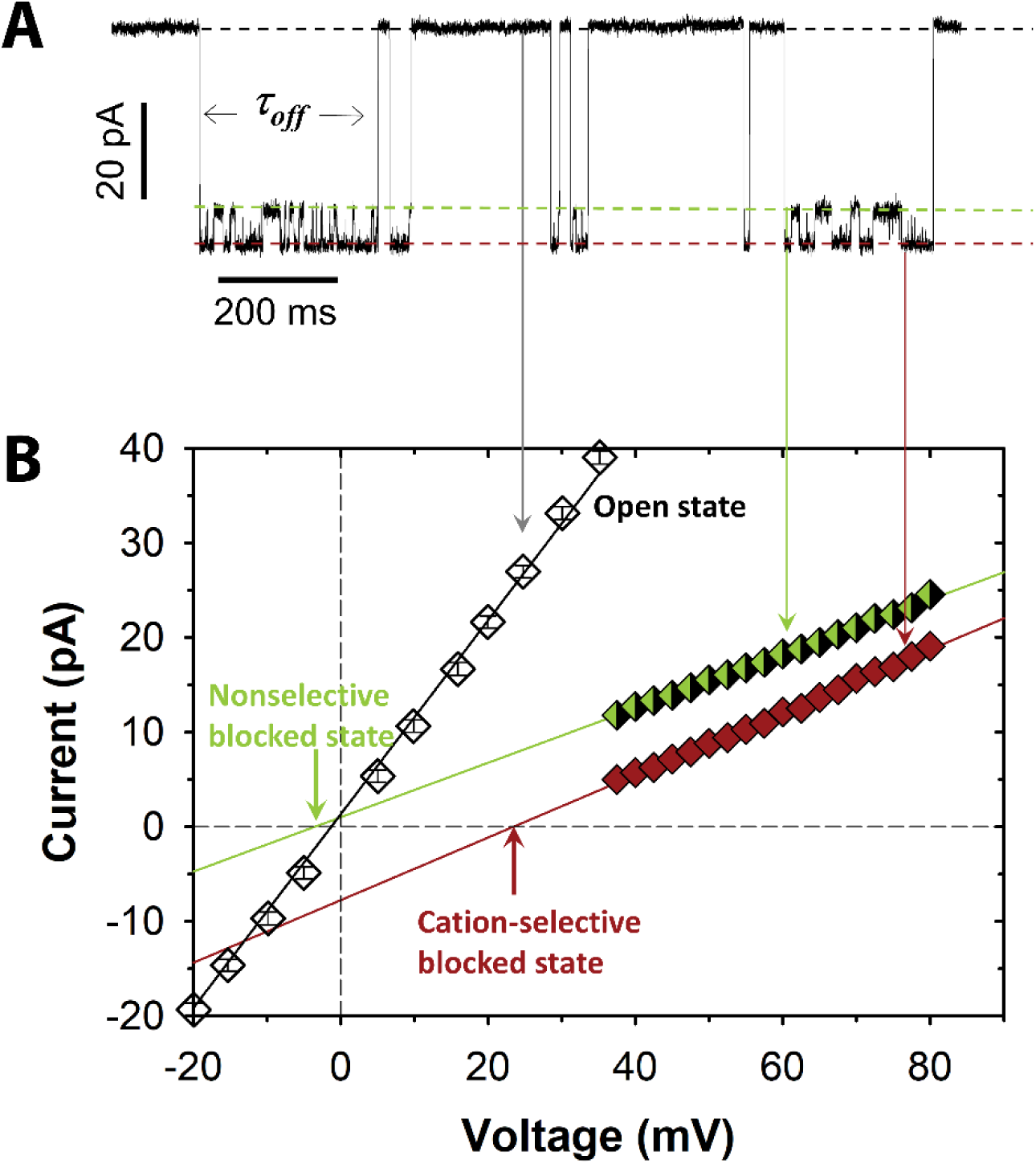
Salt gradient (0.2 M KCl *cis* / 1.0 M KCl *trans*, 5 mM HEPES, pH 7.4) imposed across the membrane with a single MspA nanopore allows for observation of dynamics of αSyn / pore interaction. (A) MspA open pore current (top level) and splitting in the residual current of the blocked state (two bottom levels). (B) The corresponding current-voltage relationship demonstrates the ion selectivity of the observed conductive states of MspA: the open state of the pore is non-selective, and its reversal potential is close to zero; the two conductive states of the αSyn blocked state are either non-selective (green diamonds; reversal potential is −3.7 ± 0.6 mV, giving a 1.2 Cl^−^/1 K^+^ ratio) or cation-selective (red diamonds; reversal potential is 23.2 ± 0.7 mV, giving a 0.23 Cl^−^ / 1 K^+^ ratio). Error bars represent 68% confidence intervals of the mean value and where not visible are smaller than the size of the data point.

Observation of the last state before the αSyn molecule is released from the pore lumen reports on the direction taken by the molecule upon its release: if the C-terminal domain is last observed, αSyn retracted from the pore to the side of capture; if the N-terminal domain is last observed, the αSyn translocated the membrane through the nanopore (14). In principle, this measurement technique can be applied to any polypeptide by attachment of a charged “selectivity tag” to its terminus; here, we exploit αSyn’s natural structure that already contains such tag domain.

Because the αSyn molecule is expected to partition into the nanopore with its polyanionic C-terminal end first (while its N-terminal is membrane-bound), one can reasonably expect that the resulting cation-selective state should be observed at the beginning of each blockage event. This is indeed true, as shown in Fig. 4, where both types of blockage events are shown to be initiated in the cation-selective state.

**Figure 4.**
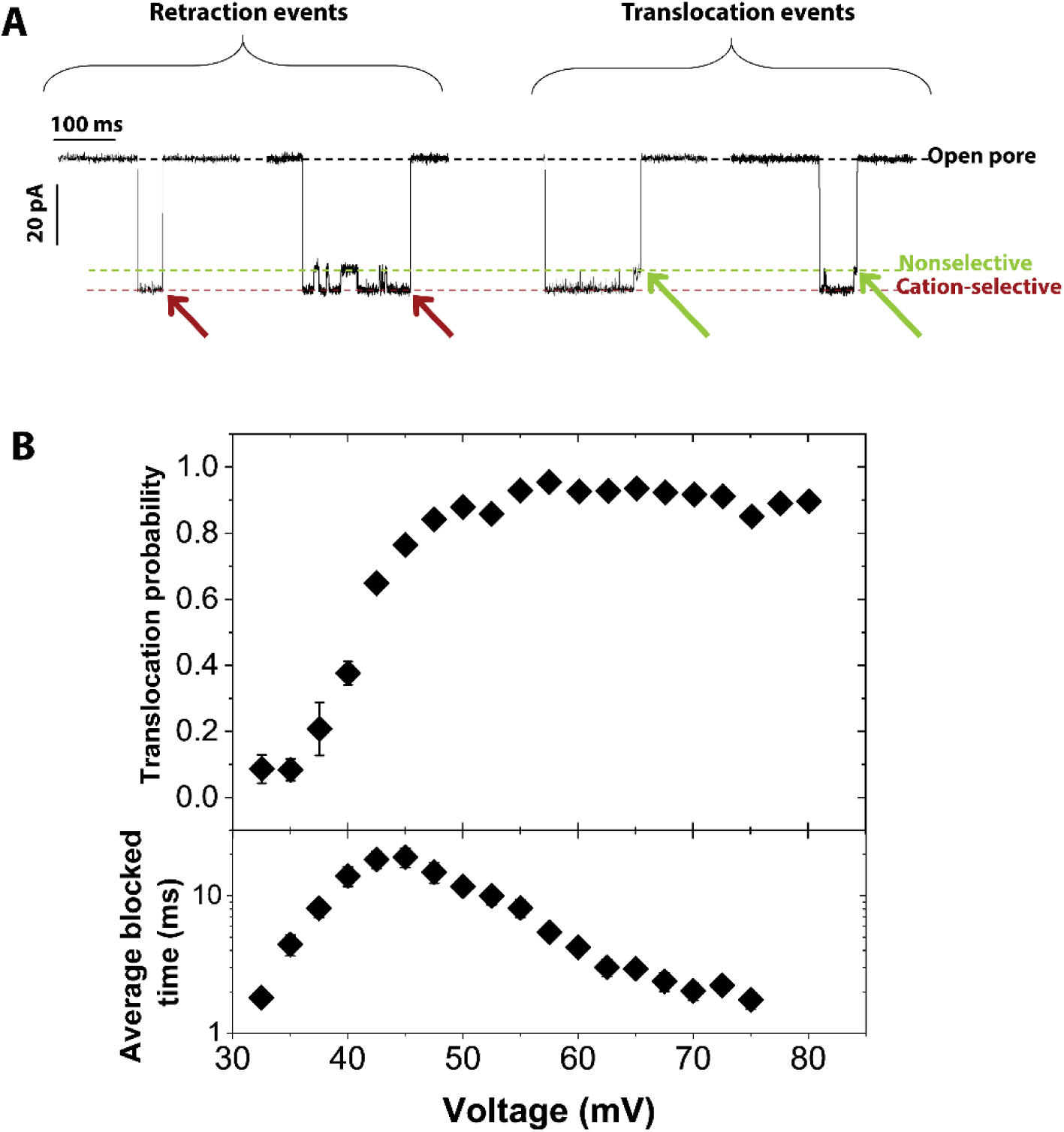
(A) Sample events demonstrating the technique of classifying retraction vs. translocation events of αSyn interaction with MspA nanopore. The salt gradient is 0.2 M /1.0 M KCl, applied voltage is 45 mV. Two levels within the residual current of the blocked state are clearly visible; the deeper, cation-selective state is indicative of the presence of the C-terminal domain in the pore, while the shallower, non-selective state indicates the presence of the N-terminal domain. Retraction events are characterized by starting and ending from the same cation-selective state (C-terminal tail is both captured first and leaves the pore last). Translocation events start in the cation-selective state but end in the non-selective state (the C-terminal domain is captured initially, but the N-terminal domain is last in the pore). The substate at the end of each event, which is used to classify retraction vs translocation events, is marked by an arrow. (B) Voltage dependence of the probability of αSyn translocation by MspA pore and average time of the entire blockage event, obtained in 0.2/1.0 M gradient of KCl, buffered with 5 mM HEPES, pH 7.4. Top panel presents the percentage of events in which the uncharged selectivity tag was observed at the end of the blockage event. Bottom panel gives average event durations. Error bars represent 68% confidence intervals estimated from bootstrap resampling of the observed distributions.

To rephrase, an αSyn molecule trapped by a MspA pore has only two ways out: (i) retract from the pore and end up on the side of the membrane from where it originally arrived, or (ii) unbind from the membrane surface and translocate through the pore. Depending on how deeply the αSyn molecule penetrates the pore, one should expect at least two types of retraction events. The first can be very brief and is limited only to the polyanionic C-terminal domain partitioning into the pore, followed immediately by retraction before the N-terminal domain ever enters the pore constriction. The other retraction event involves excursions of the N-terminal domain of αSyn into the pore. Trapped by the pore, the αSyn molecule can fluctuate between these states, observed experimentally as transitions between cation-selective and non-selective sub-states of the residual current. Regardless of the event type, if the cation-selective state is the last observed, the C-terminal selectivity tag was the last region of the αSyn molecule within the pore lumen. Examples of such types of retraction events are readily observed in our current records and are presented in Fig. 4A.

An electric signature of the translocation event, on the contrary, includes always at least one transition between the residual current substates: first, the C-terminal partitioning (cation-selective state) necessarily followed by moving of the nearly-net-neutral N-terminal domain into the pore (non-selective state). From that point the αSyn can clear the channel immediately or fluctuate in the pore for some time; however, the N-terminal domain of the molecule is the last one to leave the pore (Fig. 4B). We show here that the principle of following the charged domains of analyzed protein in a salt-gradient system for real-time monitoring of protein’s translocation, first demonstrated for mitochondrial VDAC and αSyn system (14) applies equally well to the structurally non-homologous proteinaceous pore formed by mycobacterial porin MspA.

## Discussion

Protein complexation involves different kinds of interactions between the participating molecules. These interactions are usually characterized as Coulombic, Van der Waals, hydrophobic, hydrogen bonding, salt-bridging, etc., though the underlying physical forces are not always easy to separate into these specific types. In any case, for membrane-bound proteins, the overall energetics of the protein-protein interaction can now be supplemented by an additional well-defined term that is distinctly different from those listed above – by the force exerted on charges of the interacting molecules by a transmembrane electric field.

In this sense, the voltage-activated association of a peripheral and integral membrane protein, such as that demonstrated previously for αSyn with VDAC or α-hemolysin, represents a novel mechanism for lipid-mediated protein-protein interactions. While the peripheral protein has two basic requirements—a membrane anchoring domain and a disordered polyionic domain (11)— the question of the requirements on the channel have remained open. Do residue-specific interactions between the channel and its polypeptide inhibitor qualitatively alter the interaction pattern? Are the anion selectivity and voltage-induced gating properties of VDAC, or other yet to be determined structural motifs or channel functionalities, required for complex formation? Strong evidence suggests otherwise. The energy landscape modeling approach (16) accurately describes a broad range of experimental data for the VDAC channel without including specific interactions. Most notably, energy landscape modeling completely elides specific interactions between structures of the channel and the captured polypeptide chain. It relies solely on mean-field concepts for hydrodynamic drag and electrical forces drawn from polynucleotide motion in nanopores (31-35). Energy landscape modeling has quantitatively described the osmotic forces acting on αSyn in VDAC under an electrolyte concentration gradient (14). It has revealed the effect of lipid composition on the conformational distributions of membrane-bound αSyn (12, 13, 36). Energy landscape modeling was shown to give the position of post-translational modification mimics (6), and, in the case of tubulin, can distinguish between α and β isoforms (15). Thus, it is expected that in the absence of channel specificity, an unrelated channel that does not gate and is not ion selective should interact with VDAC’s polypeptide inhibitors in a qualitatively similar way. Here, we have confirmed this prediction, showing that MspA can be substituted for VDAC with no qualitative differences in its interaction with αSyn. VDAC’s capture rate for αSyn is rather higher, probably due to the higher anion selectivity of the channel, but otherwise all qualitative features of the interaction—the transition from reversible blockages of the retraction regime to those of translocation and the alterations of the selectivity of the channel based on the charge density of the αSyn residues in the channel interior—remain identical.

To conclude, we point out that the multiple short- and long-range forces, described in the literature for protein complexation, now should be augmented by a physical interaction that is due to the unique properties of cellular and organelle membranes, the important structural elements of the cell architecture. A membrane’s hydrophobic core, with a thickness of a few nanometers, excludes small ions and thus provides excellent insulation, allowing building up electric fields of hundreds of kilovolts per centimeter. These large transmembrane fields govern protein-protein interactions by a mechanism that cannot occur for proteins in bulk solutions. Using the MspA nanopore reconstituted into a planar lipid bilayer we explored a new example of such membrane-catalyzed interactions. Depending on the applied voltage, the reaction equilibrium constant can vary within many orders of magnitude. Though previously studied in detail for the αSyn docking into the VDAC pore, the described mechanism remains quite new and deserves further investigations, one of which is presented in the present study.

In addition to clarifying the physics of this novel protein-protein complexation mechanism by validating the use of mean-field descriptions for the thermodynamics and kinetics of the interaction (6, 11-16), the present results also expand the suite of nanopores available for detailed study of polypeptides (37). Significant progress has already been made in this direction using aerolysin nanopores (10) and engineered nanopores (7). We now advance this approach further to allow for quantitative analysis of the fine features of protein and polypeptide binding to the membrane surface and its functional consequences for complexation. Importantly, the detailed physical model that describes the motion of a disordered polypeptide chain with a selectivity tag in the biological nanopore turns out to be quite general and thus provides for the reliable quantitative prediction of the efficiency of this type of protein-protein complexation.

## Author contributions

D. P. H., P. G., T. K. R., and S. M. B. designed the experiments; P. G. performed the experiments; P. G., D. P. H., and T. K. R. analyzed the data; D. P. H., P. G., T. K. R., and S. M.B. wrote the manuscript; A. L. and J. G. designed, produced, and supplied the MspA protein; all authors edited the manuscript.

## Acknowledgments

T.K.R. and S.M.B. were supported by the Intramural Research Program of the *Eunice Kennedy Shriver* National Institute of Child Health and Human Development of the National Institutes of Health. Certain commercial materials, equipment, and instruments are identified in this work to describe the experimental procedure as completely as possible. In no case does such an identification imply a recommendation or endorsement by the National Institute of Standards and Technology, nor does it imply that the materials, equipment, or instrument identified are necessarily the best available for the purpose.

